# Sex differences in brain metabolism assessed with whole-brain magnetic resonance spectroscopic imaging

**DOI:** 10.64898/2026.06.30.735476

**Authors:** Edgar Céléreau, Federico Lucchetti, Pascal Steullet, Zoé Schilliger, Yasser Alemán-Gómez, Raoul Jenni, Teya Petrova, Silas Forrer, Farnaz Delavari, Jean-Baptiste Ledoux, India D’Addona, Lila Wider, Maria Fernanda Rueda, Basilio Giangreco, Patric Hagmann, Kerstin Jessica Plessen, Stephan Eliez, Philippe Conus, Camille Piguet, Arnaud Merglen, Daniella Dwir, Antoine Klauser, Paul Klauser

## Abstract

Sex differences in brain disorders span age at onset, symptom profiles, disease course and treatment response, and may partly reflect underlying differences in cellular metabolism. Indeed, in vivo evidence of sex-related neurometabolic variation remains sparse, with heterogenous and conflicting findings. Using fast high-resolution whole-brain three-dimensional magnetic resonance spectroscopic imaging, we mapped five brain metabolites in three independent cohorts of healthy participants (total n = 114). In a discovery sample of adolescents scanned at 3 Tesla (3T) (n = 61), males showed higher total N-acetylaspartate (tNAA) across widespread gray matter regions. Regional analyses further revealed opposing sex patterns with a complementary higher total creatine (tCr) observed in females, motivating examination of their ratio as an integrative metabolic index. The tNAA/tCr ratio was consistently higher in males in the discovery sample and this finding was replicated across two independent young-adult samples (3T, n = 26; 7T, n = 27), with a widespread gray and white matter distribution. This tNAA/tCr ratio may link neuronal mitochondrial metabolism with cellular energy buffering, positioning it as a potential index of bioenergetic balance relevant for conditions showing both sex differences and altered neurometabolism, notably multiple sclerosis, Alzheimer disease, and psychosis. Together, these findings reveal a reproducible, distributed metabolic sexual dimorphism in the human brain, and underscore the importance of accounting for sex-specific neurometabolic profiles in studies of brain health and disease.

## Introduction

Sex differences in psychiatric and neurological disorders have been repeatedly reported, spanning age at onset, symptom profiles, comorbidity patterns, and treatment response^1,2^. This may reflect sex-specific variations in brain cellular composition and energy metabolism. Indeed, the evidence suggests that sex steroids may play a regulatory role in mitochondrial function, glucose metabolism, and cellular bioenergetics^3^. This suggests the possibility of differences in brain metabolic organization between males and females, beyond the macroscopic anatomical differences that have already been observed^4^.

Proton magnetic resonance spectroscopy (^1^H-MRS) enables the study of multiple brain metabolites in vivo, while magnetic resonance spectroscopic imaging (MRSI) combines this spectral information with spatial localization. Recent advances in proton MRSI now allow the mapping of metabolites in three dimensions (3D-MRSI) across the whole brain with high resolution in clinically feasible acquisition times^5^. Whole-brain 3D-MRSI has been shown to recover reproducible regional metabolite distributions^6^ and sufficient sensitivity to capture aspects of the underlying biochemical brain organization^7^. At 3T, 3D-MRSI reliably resolves major brain metabolites including total N-acetylaspartate (tNAA: N-Acetylaspartate [which accounts for 75-90% of the tNAA signal] + N-acetylaspartylglutamate), total creatine (tCr: Creatine + Phosphocreatine), glutamate and glutamine (combined in Glx), choline-containing compounds (Cho: Glycerophosphocholine + Phosphocholine), and myo-inositol (Ins)^8^. These metabolites provide complementary biological information: total-NAA is often considered a marker of neuronal integrity and mitochondrial metabolism; tCr reflects the creatine kinase system involved in cellular energy buffering; Glx relates to excitatory neurotransmission and glutamate-glutamine cycling; Cho to membrane turnover; and Ins to glial-associated processes.

Previous studies have explored sex difference in brain metabolism using single-voxel ^1^H-MRS (SV-MRS) and magnetic resonance spectroscopic imaging (MRSI) reporting metabolic sex differences limited to one or few regions of interest, but with inconsistent results. For example, sex-related differences were reported with increased tNAA and tCr in the occipital lobe and fronto-temporal white matter in males^9^, asymmetries of tCr levels in the frontal lobe of females^10^, and higher Glx and tCr in female hippocampus alongside higher Cho in the anterior cingulate cortex^11^.

In this study, we investigated the effects of sex on these five brain metabolites in the whole brain using a novel 3D-MRSI technique and spatially resolved analyses in a discovery sample of healthy adolescents and two independent replication samples of young adults. We hypothesized that, despite previous mixed and localized findings from ^1^H-MRS, 3D-MRSI might be more sensitive to reveal distributed brain metabolic differences between sexes.

## Methods

### Participants

Participants were drawn from three independent studies of healthy youth: a discovery sample scanned in Geneva at 3T (GE3T; Siemens Magnetom TrioTim 3T), comprising 61 adolescents from the Mindfulteen study, and two replication samples, including 26 healthy participants from the Lausanne psychosis cohort (LA3T; Siemens Magnetom PrismaFit 3T) and 27 healthy participants from the 22q11DS study in Geneva (GE7T; Siemens Magnetom Terra.X 7T). All studies were approved by local ethics committees and participants, and/or their legal guardians, provided written informed consent.

The Mindfulteen study (GE3T) is a randomized controlled crossover trial assessing the effects of a mindfulness-based intervention in adolescents from the general population. Participants were aged 13 to 15 years and were excluded if they had chronic somatic disease, significant medical conditions, psychotherapy in the previous 6 months, psychotropic medication in the previous month, or current or past psychiatric disorder, except anxiety disorder or past major depressive disorder. The Lausanne (LA3T) participants were healthy controls recruited in parallel to the Lausanne psychosis cohort and were required to be free of any lifetime psychiatric diagnosis and to have no first-degree relative with a psychosis-related disorder. The GE7T subjects are healthy participants from the 22q11DS study; participants younger than 13 years were excluded to better match GE3T and LA3T. For participants with multiple scans, the first scan with adequate whole-brain coverage and no significant motion artifacts was selected (details provided in further sections).

### MRI acquisition

For GE3T, Magnetic Resonance (MR) data were acquired on a 3T Magnetom TrioTim scanner (Siemens Healthineers, Forchheim, Germany) equipped with a 32-channel head coil at the Brain and Behavior Laboratory in Geneva. For LA3T, MR data were acquired on a 3T Magnetom PrismaFit scanner (Siemens Healthineers, Forchheim, Germany) equipped with a 32-channel head coil at the CIBM Center for Biomedical Imaging in Lausanne University Hospital. For GE7T, MR data were acquired on a 7T Magnetom Terra.X scanner (Siemens Healthineers, Forchheim, Germany) equipped with a 32-channel head coil in Geneva.

Each session included a T1-weighted magnetization-prepared rapid acquisition gradient echo sequence used for anatomical reference, tissue segmentation, and spatial normalization. In GE3T and LA3T, the T1w MPRAGE was acquired with 1 mm in-plane resolution and 1.2 mm slice thickness (coverage 240 × 256 × 160 voxels), with TR = 2300 ms, TE = 2.98 ms, TI = 900 ms. In GE7T, the T1-weighted MP2RAGE was acquired with different parameters due to the 7T field: 0.6mm in-plane resolution and 0.66mm thickness (coverage 396 x 416 x 256 voxels), with TR= 6000 ms, TE= 2.07 ms and TI= 800 ms.

### 3D-MRSI acquisition

Whole-brain metabolic imaging used a short-TE, compressed-sensing accelerated 3D ¹H-FID-MRSI sequence.

For GE3T, the specific sequence was a 3D ^1^H-CS-SENSE-LR-FID-MRSI sequence, described in Klauser et al^12^. Acquisition parameters were TE = 1.5 ms, TR = 372 ms, flip angle = 35°, FOV = 210 × 160 × 105 mm^3^, slab thickness 95 mm, spatial resolution 5 × 5 × 5.3 mm^3^, bandwidth 2 kHz, and 512 FID points; the water reference used TE = 1.5 ms, TR = 36 ms, and a flip angle = 3°, with a resolution of 6.6 x 6.7 x 6.6 mm^3^. The total acquisition time was 22min, including 2min for water reference acquisition.

The LA3T was acquired with 2 specific sequences that ran consecutively on the same MRI scanner. From 2019 to 2022, acquisition sequence was the same as for GE3T, with 13 subjects (50%) scanned. From 2025 to 2026, acquisition sequence was a 3D 1H-ECCENTRIC-FID-MRSI sequence, described in Klauser et al^5^, with also 13 subjects (50%) scanned. Acquisition parameters for the first were TE = 1.00 ms, TR = 353 ms, flip angle = 40°, with matching spatial resolution, bandwidth, and vector size relative to GE3T. A separate water reference was acquired (same TE; TR = 25 ms; flip angle = 3°; resolution of 6.6 × 6.7 × 6.6 mm^3^). Total acquisition time was 22 min including 2 min for water reference. Acquisition parameters for the second LA3T acquisition were TE = 0.78 ms, TR = 457 ms, flip angle = 45°, FOV = 220 x 220 x 130 mm³, slab thickness 95mm spatial resolution 5 × 5 × 5.2 mm^3^, bandwidth 1.32 kHz, and 512 FID points; the water reference used TE = 0.72 ms, TR = 460 ms, and a flip angle = 45°, with a resolution of 10 × 10 × 10 mm^3^. Total acquisition time was 8:15min, with 6:54min for MRSI and 1:21 for water reference.

The GE7T used the same ECCENTRIC-FID-MRSI sequence as the more recent LA3T. Acquisition parameters were TE = 0.68ms, TR = 400 ms, flip angle = 35°, FOV = 220 x 220 x 110 mm^3^, slab thickness of 100mm, spatial resolution 3.4 x 3.4 x 3.5 mm^3^, bandwidth 2280 Hz, and a 688 FID points. A separate water reference was acquired (TE = 0.59; TR = 404 ms; flip angle = 35°; resolution of 10 × 10 × 10 mm^3^). Total acquisition time was 11:52min, including 59sec for water acquisition.

Detailed Minimum Reporting Standards (MRSinMRS) are summarized in the Table S1.

### Reconstruction and spectral quantification

Reconstruction was performed using a low-rank model constrained by total generalized variation (TGV) with simultaneous removal of subcutaneous lipid contamination and residual water signals, following Klauser et al.^5^. After reconstruction, spatio-spectral data were quantified voxel-wise using LCModel^13^, using the separate water acquisition as the concentration reference (see Fig. S1 for spectra examples in each cohort).

The LCModel basis set included (non-exhaustive) N-Acetylaspartate (NAA), N-Acetylaspartyl-Glutamate (NAAG), Creatine (Cr), Phosphocreatine (PCr), Glycerophosphocholine (GPC), Phosphocholine (PCh), Myo-inositol (Ins), Scyllo-inositol, Glutamate (Glu), Glutamine (Gln), Lactate, GABA, Glutathione (GSH), Taurine, Aspartate, and Alanine; due to spectral overlap, analyses focused on five robustly-resolved signals for the 3T acquisitions: tNAA (NAA+NAAG), tCr (Cr+PCr), Cho (GPC+PCh), Ins (myo-inositol), and Glx (Glu+Gln). For the 7T acquisition, nine robustly resolved metabolites are analyzed : NAA, NAAG, tCr (Cr+PCr), Cho (GPC+PCh), Ins, Glu, Gln, GABA, GSH. However, in order use the 7T data as a reproduction sample of the 3T discovery sample, NAA and NAAG were used as a tNAA (NAA+NAAG) map.

Metabolite levels are reported in institutional units (IU). LCModel-derived voxel-wise quality metrics included Signal-to-Noise Ratio (SNR), Full Width at Half Maximum (FWHM), and Cramer-Rao Lower Bound (CRLB).

### Quality control and quality masks (Qmasks)

For each subject and metabolite, a binary quality mask (Qmask) was generated by combining the LCModel quality maps (SNR, FWHM, and CRLB). Voxels were labeled reliable (value = 1) if they simultaneously satisfied CRLB < 20, FWHM < 0.1, and SNR > 4; otherwise, they were marked unreliable (value = 0). Metabolic maps were masked using the corresponding Qmasks. Consistent with the source workflow, acquisitions with major artifacts (e.g., motion inferred from anatomical inconsistencies in metabolic maps and/or insufficient brain coverage as reflected by Qmasks) were excluded based on visual quality control. If a subject had multiple scans, only the first one without major artifacts was included.

Consequently, in the GE3T sample, 69 subjects were scanned, 68 of them having 2 or 3 scans at different time points, totalizing 167 3D-MRSI acquisitions. 53 acquisitions were excluded for bad quality. This selection brought the number of individual subjects with a usable 3D-MRSI scan to 61.

In the LA3T sample, 29 subjects were scanned, 5 of them having scans at multiple time points, totalizing 36 MRSI brain scans. 5 acquisitions were excluded for bad quality, bringing the number of individual subjects with a usable 3D-MRSI scan to 26.

In the GE7T sample, 32 subjects were scanned once, 4 were excluded for bad quality and 1 was excluded for a medical condition discovered during assessment. This brought the number of individual subjects with a usable 3D-MRSI scan to 27.

### Preprocessing: filtering, co-registration, partial-volume correction, normalization

The voxel-based preprocessing pipeline was designed to normalize each metabolic volume to standard space prior to voxel-wise analyses and comprised: (i) spatial filtering of spikes, (ii) partial-volume effect (PVE) correction, (iii) co-registration to T1-weighted anatomy, and (iv) warping to MNI152 space. Detailed information can be found in the original publication of this preprocessing method^6^.

Outlier “spikes” (often arising from LCModel quantification errors when the water reference is underestimated) were detected using a threshold set at the 99th upper percentile of intensities within the brain mask, then corrected via localized inpainting using a 3×3×3 median filter. Voxels with NaNs or zeros within the brain mask were repaired using biharmonic inpainting followed by localized median filtering. Filtered maps were then smoothed using a 5-mm (or 3mm for the 7T acquisitions) FWHM Gaussian kernel (chosen to match the intrinsic ∼5 mm [or 3mm for 7T] acquisition resolution).

Filtered metabolic maps were co-registered to skull-stripped (with hd-bet^14^) T1w images, using tCr as the registration reference and ANTs^15^ v2.4.1 with a rigid transform (mutual information) followed by symmetric diffeomorphic deformation (cross-correlation). The tCr map to T1w transformation is then applied to the four remaining metabolic maps to ensure all maps have similar registration.

T1w images were segmented with CAT12^16^ v12.9 to generate GM/WM/CSF probability maps in T1w space. The inverse MRSI-to-T1w transform was applied to tissue maps, and metabolite maps were corrected for GM/WM/CSF partial-volume effects using a region-based voxel-wise correction, as implemented in PETPVC toolbox^17^.

Finally, T1w images were normalized to MNI15212 using ANTs (rigid + affine with mutual information; symmetric diffeomorphic with cross-correlation), and the resulting transforms were applied to PVE-corrected metabolite maps. Normalized metabolic maps were resampled to 1×1×1 mm³ using tri-linear interpolation at each transformation stage.

All Qmasks underwent the same spatial transforms as their corresponding metabolic volumes using tri-linear interpolation.

### Derived maps (tNAA / tCr ratio)

To facilitate comparisons and reduce dependence on a single internal reference, tNAA / tCr ratio maps were computed by dividing each normalized tNAA map voxel-wise by the corresponding tCr map, adding a small constant (ε = 1×10⁻⁶) to the denominator for numerical stability; ratio computations are restricted to the brain mask.

### Voxel-wise statistical analyses

Voxel-based analyses were performed on normalized metabolic maps using a generalized linear model and FSL randomise v6.0.7.18^18^. Family-wise error correction used Threshold-Free Cluster Enhancement (TFCE^19^) with permutation testing (10,000 permutations). Poor-quality voxels were handled using the lesion-masking option in randomise: each subject’s Qmask served as a voxel-wise regressor so that unreliable voxels were excluded from the inference. Analyses were restricted to the gray and white matter, with the cerebellum and the ventricules excluded, following the Harvard-Oxford atlas^20^.

Models included sex as the primary variable of interest, with age as covariate. Total intracranial volume (TIV), which was estimated via CAT12 during the T1w processing, was included as an additional covariate in additional analyses.

### Mean extraction by region

Brain masks in standard space were generated using the Harvard-Oxford atlas: frontal, parietal, temporal and occipital lobes, with subdivisions for gray and white matter as well as thalamus, putamen, pallidum and caudate; each region was also subdivided between left and right side. A global quality mask was generated for GE3T by multiplying all individual Qmasks together of the subjects in the GE3T sample. This generated a global Qmask per metabolite. Each regional brain mask (e.g., frontal gray matter) was then multiplied by this global Qmask to extract reliable values for each sample in each brain region. The mean value for each subject was extracted from each of these brain regions.

Mean values per sex were plotted in R v. 4.5.2 using ggplot2^21^and ggridges^22^ packages.

### Code availability

The preprocessing (https://github.com/MRSI-Psychosis-UP/mrsiprep) and voxel-based (https://github.com/MRSI-Psychosis-UP/VLAD) pipelines are publicly available on GitHub.

## Results

Demographics for the 3 samples are displayed in Table 1. Sex distribution was balanced and similar across samples (Fisher’s p = 0.762). Age was not different between males and females inside each sample but differed between samples (p<0.001), allowing to cover a range from puberty (∼13) to adulthood (∼35). Total Intracranial Volume (TIV) was larger in males than in females, and similar in the 3 samples despite age difference (p = 0.754).

**Table 1:** Demographics data for the 3 samples (GE3T, LA3T and GE7T)

|  | **Geneva 3T (n=61)** | **Lausanne 3T (n=26)** | **Geneva 7T (n=27)** |
| --- | --- | --- | --- |
| **Sex** |  |  |  |
| No. Females (%) | 35/61 (57%) | 13/26 (50%) | 16/27 (59%) |
| No. Males (%) | 26/61 (43%) | 13/26 (50%) | 11/27 (41%) |
| **Age, mean (SD)** | 14.56 (0.92) | 26.24 (7.54) | 18.83 (4.47) |
| Females | 14.59 (0.93) | 28.18 (9.65) | 19.36 (4.39) |
| Males | 14.50 (0.92) | 24.30 (4.12) | 18.07 (4.68) |
| p-value | 0.702 | 0.201 | 0.477 |
| **Total Intracranial Volume, mean (SD), cm<sup>3</sup>** | 1487 (104) | 1507 (124) | 1488 (140) |
| Females | 1452 (102) | 1442 (97) | 1425 (115) |
| Males | 1534 (88) | 1572 (115) | 1579 (125) |
| p-value | 0.001 | 0.005 | 0.004 |

In the discovery sample (GE3T), VBA revealed higher tNAA levels in males compared with females, forming a widespread cluster encompassing mostly gray matter (Fig. 1A-B). This effect remained unchanged after adjustment for TIV (Fig. S2). No other metabolite showed significant voxel-wise differences between sexes. Regional summaries (Fig. 1C) showed a consistent pattern of higher tNAA in males and higher tCr in females across brain regions.

**Fig. 1.**
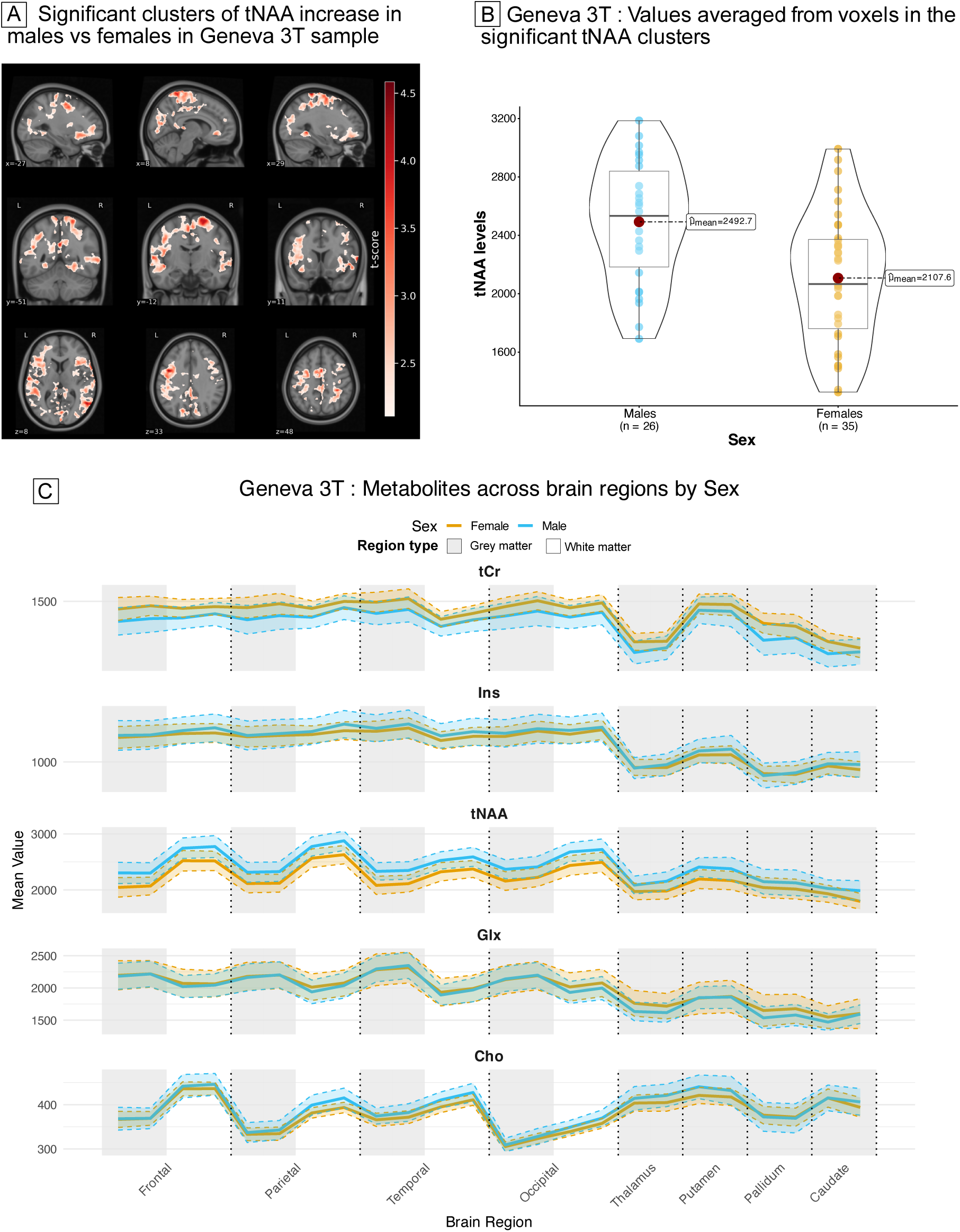
Absolute metabolite differences in the discovery sample (GE3T). (A) Voxel-based analysis of tNAA showing clusters with higher values in males compared with females (TFCE-corrected p < .05). (B) Individual mean tNAA values extracted from significant clusters in the GE3T sample. (C) Regional mean concentrations of the five metabolites by sex. Although not statistically significant, tNAA shows a consistent elevation in males, whereas tCr shows a consistent elevation in females; no consistent pattern is observed for other metabolites.

Given these opposing trends, we next examined the tNAA / tCr ratio as an index integrating complementary aspects of brain metabolism and a more standardized way to compare the samples. The tNAA / tCr ratio was higher in males compared to females in GE3T sample (Fig. 2A-B) and in the two other samples (LA3T [Fig. 2C-D], GE7T [Fig. 2E-F]) showing a similarly widespread pattern across both gray and white matter.

**Fig. 2.**
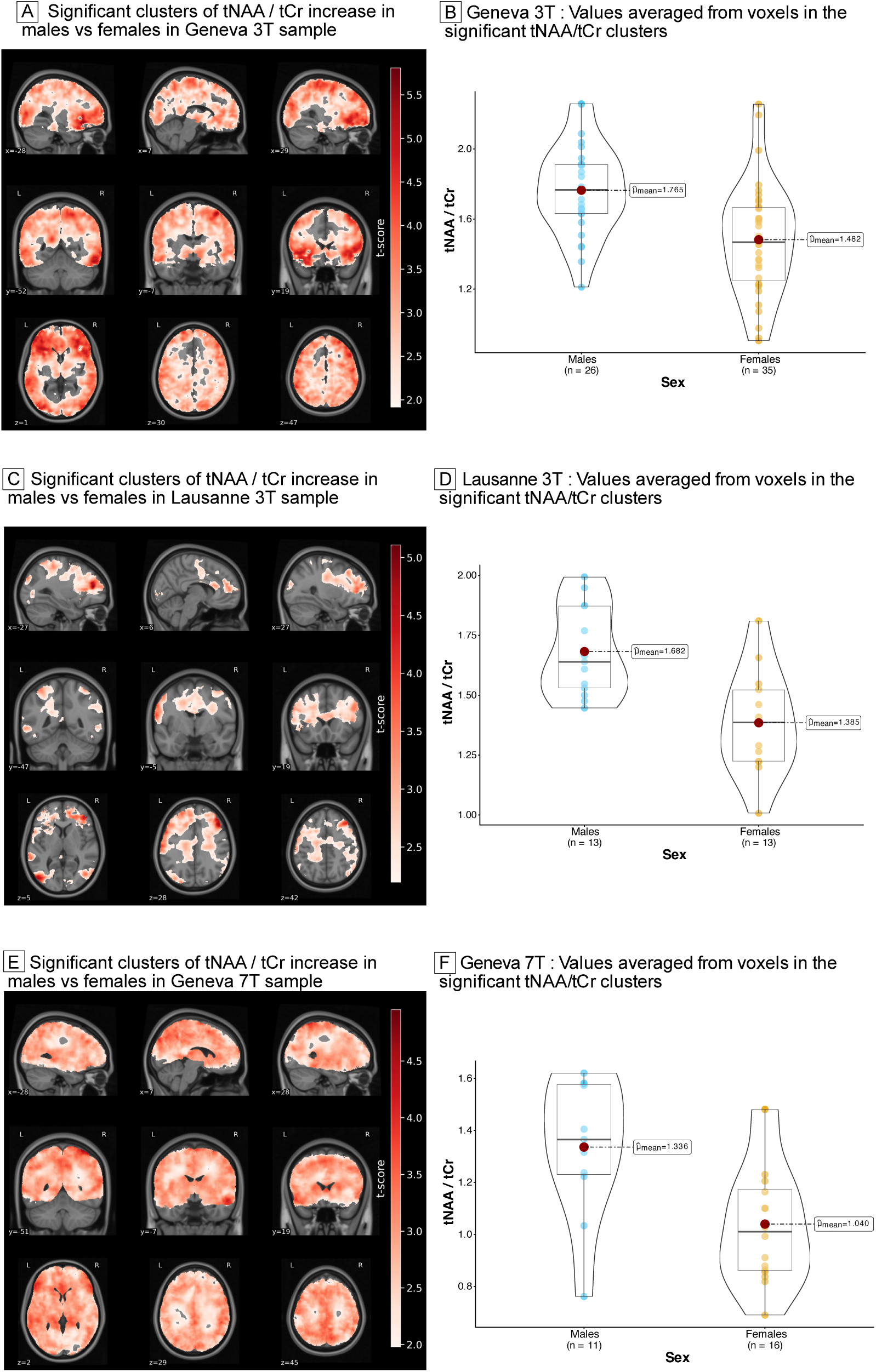
Voxel-based analyses of tNAA/tCr ratio across three independent samples. (A, B) GE3T (discovery sample): (A) clusters with higher tNAA/tCr in males vs females (TFCE-corrected p < .05); (B) corresponding individual mean values extracted from significant clusters. (C, D) LA3T (replication sample): same analyses as in panels A–B. (E, F) GE7T (replication sample): same analyses as in panels A–B.

## Discussion

To the best of our knowledge, this is the first in vivo study showing replicable widespread metabolic sex differences in the brain. While tNAA levels were higher in males in the discovery sample, the tNAA / tCr ratio showed a consistent increase in males across 3 independent samples. The spatial extent of this effect, involving both gray and white matter, supports the presence of a distributed metabolic dimorphism rather than a localized regional difference. These findings suggest that sex is associated with broad differences in the cerebral metabolic organization detectable in vivo.

The biological interpretation of this finding is supported by the complementary roles of tNAA and tCr in brain energy metabolism. Beyond its widespread use as a marker of neuronal integrity, NAA has been implicated in neuronal osmoregulation, axon-glial signaling, and the transfer of acetate units required for lipid synthesis. NAA is synthesized in neuronal mitochondria from acetyl-CoA and aspartate by the enzyme NAT8L^23^. While acetyl-CoA is the initial fuel for the tricarboxylic acid (TCA) cycle, aspartate is tightly coupled with the malate-aspartate shuttle and derived from oxaloacetate through transamination reactions that depend on the TCA cycle flux^23^.

Through these pathways, NAA synthesis is tightly connected to neuronal mitochondrial activity, glucose uptake and substrate availability, redox balance, and oxidative metabolism. These pathways are known to be modulated by sex hormones^24^ and therefore this increase in tNAA in males may reflect a sex-related difference in neuronal mitochondrial metabolism.

The increase in tNAA levels in males also encompassed white matter, suggesting differences that may extend to neuronal cell bodies. Indeed, NAA provides acetate to oligodendrocytes for myelin lipid synthesis and maintenance, thereby linking neuronal mitochondrial metabolism to axonal integrity and myelination^25^. The tNAA signal also includes NAAG, one of the most abundant neuropeptides in the brain. NAAG is particularly present and detected in the white matter in the human brain in vivo^26^. It acts mainly through metabotropic glutamate receptor 3, where it modulates pre-synaptic glutamate release and modulates excitatory neurotransmission^27^.

On the other hand, tCr reflects the creatine kinase system^28^, which buffers and rapidly regenerates ATP in the cytosol to meet fluctuating energetic demands and is therefore central to short-term cellular energy homeostasis. In the brain, creatine is also involved in osmoregulation, particularly in astrocytes, and emerging evidence suggests that it may have neuromodulatory properties, including interactions with GABAergic signaling^28,29^. Although this neurotransmitter-like role remains less established than its energetic function, it reinforces the view that tCr should not be considered only as a passive normalization signal, but as a biologically meaningful marker of energy buffering and cellular homeostasis.

The shift in the balance between tNAA and tCr could therefore indicate a differentiated bioenergetic response depending on sex, potentially reflecting complementary strategies in meeting energy demands: relatively greater sustained ATP production via mitochondrial pathways in males reflected by increased tNAA or relatively greater creatine-dependent buffering of acute energetic needs in females reflected by increased tCr. A substantial body of research has demonstrated that sex hormones, notably estrogen and progesterone, have the capacity to regulate various physiological processes, including glucose metabolism, oxidative phosphorylation, respiratory-chain activity, and oxidative stress responses^30,31^. In the context of chronic stress or psychiatric conditions, such mechanisms may hold particular relevance, as evidenced by the documented differences in mitochondrial responses to glucocorticoids and estrogens^25^. Finally, because both NAAG and creatine may influence neurotransmission, the observed tNAA/tCr difference may also reflect sex-related variation in the coupling between energy metabolism and excitatory/inhibitory signaling.

Therefore, the observed sex difference may have clinical relevance because several neurological and psychiatric disorders show both sex differences and alterations in NAA-related metabolism. Multiple sclerosis (MS), which is more prevalent in women but often progresses more severely in men^32^, is characterized by reduced NAA levels, a common marker for axonal injury and neurodegeneration. Recent evidence suggests that oxidative damage to NAT8L may contribute to impaired NAA synthesis, thus resulting in neurodegeneration^33^. Creatine metabolism may also be relevant in MS, as altered creatine levels in the nervous system have been described in human studies, with impaired creatine homeostasis proposed to reflect a low metabolic state of the brain in this demyelinating disease^34^. Differences in tNAA/tCr could therefore be relevant for interpreting metabolic vulnerability and neurodegenerative trajectories in a sex-stratified strategy.

Alzheimer disease provides another important clinical context. Women have a higher lifetime risk of Alzheimer disease, and sex-dependent mechanisms involving mitochondrial function, inflammation, lipid biology, and hormonal regulation are increasingly recognized^35^. SV-MRS studies in Alzheimer disease commonly report reduced NAA, often alongside increased myo-inositol, reflecting neuronal injury and glial-associated processes^36^. Whole-brain MRSI studies have further shown distributed neurometabolic abnormalities in prodromal and dementia stages of Alzheimer disease^37^. Phosphorus MRS also revealed that women at risk for Alzheimer disease may exhibit an altered creatine kinase equilibrium and elevated ATP utilization after menopause in disease-related vulnerable regions^38^. Together, these findings suggest that sex-related variation in tNAA/tCr may capture a biologically relevant axis in Alzheimer disease, linking neuronal metabolic integrity with creatine-dependent energetic buffering in a context where female endocrine aging may reshape cerebral bioenergetic vulnerability.

Psychosis also shows sex differences in onset, course, symptom dimensions and recovery^39^. Converging evidence points towards a possible mitochondrial impairment, altered glucose utilization and metabolic reprogramming^40^, while reduced tNAA is one of the most reproducible neuroimaging alterations found in chronic psychosis^41^. Creatine-related pathways may also be relevant in psychosis: phosphorus MRS studies have reported reduced creatine kinase reaction rates in first-episode psychosis, suggesting that abnormalities in energetic buffering may be present early in the illness^42^. Sex variation in tNAA/tCr may therefore contribute to this heterogeneity and could moderate the expression of metabolic alterations across illness stages. Together, these findings suggest that sex-related variation in tNAA and tCr metabolism may represent a transdiagnostic mechanism with potential clinical relevance.

This study has limitations, notably the modest size of the replication samples and the absence of menstrual cycle assessment, which limits biological interpretation. Metabolites measured via 3D-MRSI also remain partially nonspecific, thus reflecting composite processes rather than isolated pathways. Nevertheless, the novel application of voxel-based analysis to whole-brain 3D-MRSI and the replication of our results in three independent cohorts is a major strength pushing towards sex-specific considerations when studying brain disorders.

## Conclusion

The sensitivity of the 3D-MRSI revealed a sex difference of the tNAA / tCr ratio in the whole-brain of healthy participants, with an increase of this ratio in males compared to females in 3 independent samples. Although the biological mechanisms of these findings remain to be confirmed in vivo, they highlight the need to consider metabolic differences between sexes in health and disease.

## Acknowledgments

We gratefully acknowledge the entire Mindfulteen team who contributed to the acquisition of the 3D-MRSI dataset for Geneva 3T, along with all participants; all the participants of the Lausanne 3T dataset; and all the team of the 22q11.DS cohort in Geneva, along with the participants, for the Geneva 7T dataset.

