## Supplementary Material for "Sex differences in brain metabolism assessed with whole-brain magnetic resonance spectroscopic imaging"

**Content**

Figure S1: Spectra example of the 3 cohorts

Figure S2: Voxel-based analysis of tNAA with TIV as cofactor

Table S1: MRSinMRS table (MRSI acquisition parameters)

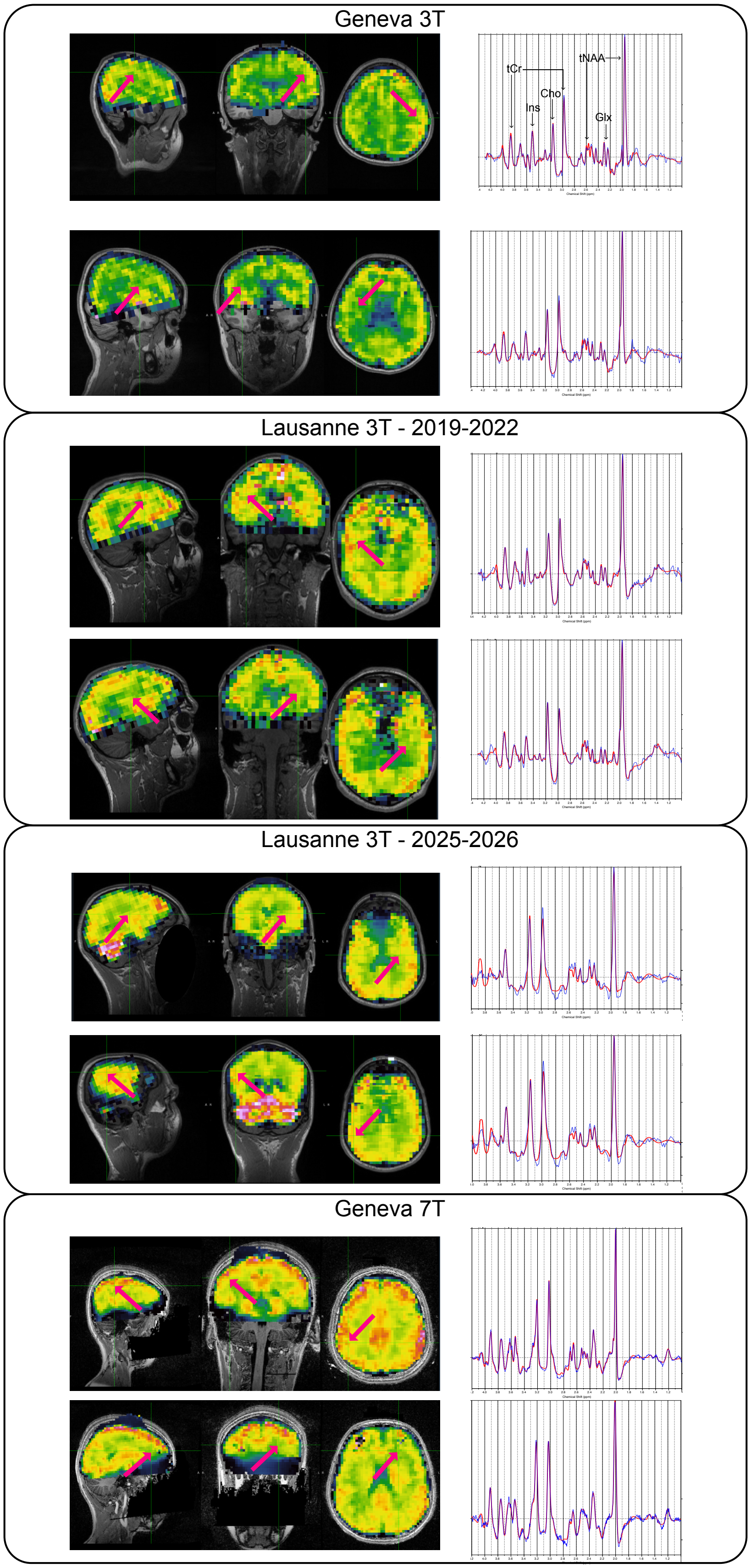


*Fig S1: Spectra example showing one spectrum in the GM and one in the WM for each acquisition (GE3T, LA3T, GE7T)*

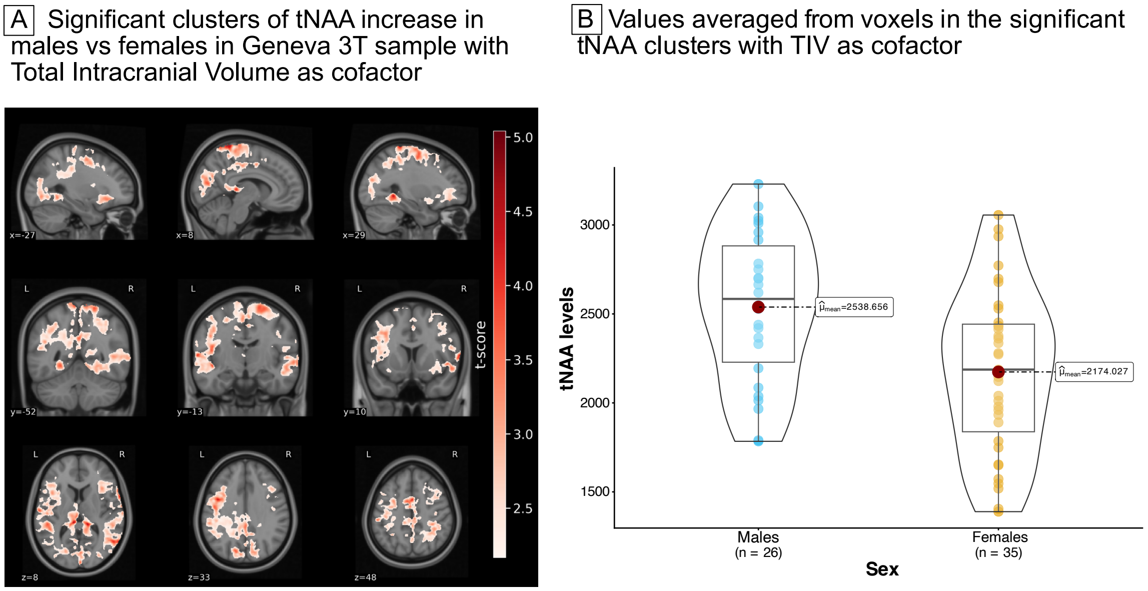


*Fig. S2: Voxel-based analysis of tNAA, with TIV as cofactor of the analysis (in addition to age and sex) showing clusters with higher values in males compared with females (TFCE-corrected $p < .05$).*

### STable 1: MRSinMRS information

| Category | Geneva. 3T | Lausanne. 3T (2019-2022) | Lausanne. 3T (2025-2026) | Geneva. 7T |
|---|---|---|---|---|
| Scanner | 3T Magnetom TrioTim (Siemens) | 3T Magnetom Prisma Fit (Siemens) | | 7T Magnetom Terra.X (Siemens) |
| RF coils | 32Rx ch 1H head coil | 32Rx ch 1H head coil | | 32Rx ch, 8Tx ch, 1H head coil |
| Coil elements | HEA;HEP | HEA;HEP | | ALL |
| Sequence | 3D 1H-FID-MRSI (CS-accelerated) | | 3D ECCENTRIC 1H-FID-MRSI | |
| Orientation | Transverse | | | |
| Rotation (deg) | -2 | 0 | -0.79 | -6.13 |
| TE (ms) | 1.5 | 1.0 | 0.78 | 0.68 |
| TR (ms) | 372 | 353 | 457 | 400 |
| Averages | 1 | 1 | 1 | 1 |
| Flip angle (°) | 35 | 40 | 45 | 35 |
| FOV (mm) | 210 × 160 × 105 | 210 × 160 × 95 | 220 x 220 x 130 | 220 x 220 x 110 |
| Slab thickness (mm) | 95 | 95 | 95 | 100 |
| Slabs | 1 | 1 | 1 | 1 |
| Resolution ($mm^3$) | 5 × 5 × 5.3 | 5 × 5 × 5.3 | 5 x 5 x 5.2 | 3.4 x 3.4 x 3.5 |
| Spectral bandwidth (Hz) | 2000 | 2000 | 1320 | 2280 |
| FID points / Vector size | 512 | 512 | 512 | 688 |

| Category | Geneva. 3T | Lausanne. 3T (2019-2022) | Lausanne. 3T (2025-2026) | Geneva. 7T |
|---|---|---|---|---|
| Acquisition duration (ms) | 256 | 256 | 389 | 302 |
| Matrix size | 42 × 32 × 20 | 42 x 32 × 20 | 44 x 44 x 25 | 64 x 64 x 31 |
| Water reference TE (ms) | 1.5 | 1.07 | 0.72 | 0.59 |
| Water reference TR (ms) | 36 | 25 | 460 | 404 |
| Water reference flip angle (°) | 3 | 5 | 45 | 35 |
| Water reference resolution ($mm^3$) | 6.6 × 6.7 × 6.6 | 6.6 × 6.7 × 6.6 | 10.0 x 10.0 x 10.0 | 10.0 x 10.0 x 10.0 |
| Water reference FID points | 16 | 16 | 512 | 688 |
| Averaging mode | N.A. | | | |
| Water suppr. | WET water suppr. | | | |
| Water suppr. BW (Hz) | 60 | 60 | 80 | 120 |
| Spectral suppr. | None | | | |
| Measurements | 1 | 1 | 1 | 1 |
| Saturation bands | 2 bands, 20 mm thickness | 2 bands, 20 mm thickness | 1 band, 20mm thickness | 2 bands, 30mm thickness |

| Category | Geneva. 3T | Lausanne. 3T (2019-2022) | Lausanne. 3T (2025-2026) | Geneva. 7T |
|---|---|---|---|---|
| Compressed sensing | Acceleration factor 3.3 | | Acceleration factor 2.5 | |
| Preparation scans | 4 | 4 | 4 | 4 |
| Dimension | 3D | | | |
| Delta frequency (ppm) | 0.00 | 0.00 | 0.00 | -2.7 |
| Phase encoding | Elliptical | | | |
| Remove oversampling | N.A. | | | |
| Shim mode | Advanced | | | |
| Data processing | Low-rank + TGV reconstruction; lipid/water removal | | Low-rank + TGV reconstruction for ECCENTRIC; lipid/water removal | |
| Quantification | LCModel | | | |
| Metabolite basis set (LCModel) | NAA, NAAG, Cr, PCr, GPC, PCh, mI, sI, Glu, Gln, Lac, GABA, GSH, Tau, Asp, Ala | | | |
| Combined metabolites | tNAA (NAA+NAAG), tCr (Cr+PCr), Cho (GPC+PCh), Ins (mI), Glx (Glu+Gln) | | | NAA, NAAG, tCr (Cr+PCr), Cho (GPC+PCh), Ins, Glu, Gln, GABA, GSH |
| Quality metrics | SNR, CRLB, FWHM | | | |
